# Loss of Hippo signaling causes transdifferentiation of neural retina between the optic fissure edges causing coloboma

**DOI:** 10.64898/2026.03.13.711620

**Authors:** Uma M. Neelathi, Daniel Sanchez-Mendoza, Sydney Steele, Rachel Aguda, Brian. P. Brooks

## Abstract

Optic fissure (OF) is a transient structure in the ventral optic cup, which acts as a conduit for periocular mesenchyme cells to enter the eye, forming hyaloid vasculature and retinal ganglion axons to exit. Optic fissure closes to form a continuous layer of retinal pigment epithelium and neural retina. Failure of OF closure results in coloboma, which is mostly genetic in nature. The severity of blindness depends on the tissue it effects and accounts for 10% of childhood blindness. In the current study, we describe coloboma pathogenesis caused by hippo effectors *yap1* and *wwtr1.* Both the paralogs are expressed in the OF edges, possibly in the pioneer cells. *wwtr1* homozygotes do not have coloboma, while *yap1* homozygotes have coloboma and pigment defects which are exacerbated by absence of one copy of *wwtr1* (*yap1^-/-^; wwtr1^+/-^*). The coloboma observed in these mutants is not due to defective optic cup morphogenesis nor an overgrown optic nerve. The pigment defects are more pronounced at the OF with complete absence of RPE specific transcription factors *mitf*A, *tfec,* and pigmentation gene *dct*. On the other hand, NR specific genes are upregulated and the unpigmented region at the OF have transdifferentiated retinal ganglion cells, amacrine, and photoreceptor cells. Our observations indicate that in the absence of *yap1* and *wwtr1*, the cells at the OF cannot attain a conducive state to fuse nor they maintain the RPE specific fate and instead they transdifferentiate into unpigmented retina, causing a steric block for fusion, resulting in coloboma.

## Introduction

Development of the vertebrate eye is an orchestrated process, which is coordinated between neuroectoderm, surface ectoderm and extra-ocular mesenchyme. The optic primordia evaginate from the neuroepithelia of the ventral forebrain as an optic vesicle and elongates to approach the surface ectoderm. There it invaginates to form a bi-layered optic cup, coincident with the surface ectoderm invaginating as the lens vesicle^1^. Optic cup morphogenesis is considered complete when the lens pinches off from the surface ectoderm^2^. The inner layer of the optic cup forms the neural retina (NR); in zebrafish, a patch of cells in the dorso-medial portion of the outer layer forms the retinal pigment epithelium (RPE). The remainder of the cells in the outer layer undergo rim movements to integrate into the inner layer to form NR^3–5^. Morphogenesis of OC, results in the formation of a ventral fissure called optic fissure (OF), which extends through the optic stalk (OS) and OC. This fissure acts as a channel for the periocular mesenchyme (POM) cells to enter the eye to form hyaloid vasculature and the retinal ganglion cell axons to exit the eye^6^. The OF has its own micro-niche of cell diversity comprising of the RPE, NR, and POM, which is itself comprised of cranial neural crest cells (NCC) originating form neural plate border and endothelial cells from mesoderm^7^. Signaling between the NR, RPE and POM results in the breakdown of the basement membrane surrounding the OF edges^8^. The cells lining the optic fissure, appose and fuse—termed “pioneer cells”--undergo morphologic changes and a partial epithelial to mesenchymal transition via differential expression of cadherins^9,10^.

Failure of OF edges to fuse results in coloboma, which occurs in 1 in 10,000 live births and accounts for approximately 10% of childhood cases^11,12^. Coloboma can affect any/all the tissues in the proximal-to-distal axis (i.e., the iris, retina, macula and/or optic nerve) with the degree of macular involvement largely determining visual acuity^11^. Coloboma can be majorly classified into two groups: 1) morphogenetic, due to defects in the formation of the optic cup; 2) a primary failure of fusion, despite normal morphogenesis and approximation of the edges of the OF^9^. Coloboma may be non-syndromic or syndromic (associated with abnormalities in other organ systems)^13^. Although environmental and dietary exposures may play a role in coloboma pathogenesis^14^, population-based studies suggest a strong genetic component^15–17^. Several genes and multiple signaling systems have been identified as causes of both syndromic and non-syndromic coloboma^18,19^.

Hippo signaling is an evolutionary conserved, serine/threonine kinase cascade which plays an important role in control of organ size, apoptosis, cell proliferation and cell shape^20,21^. *YAP1* and *WWTR1* are the effector molecules of Hippo signaling, working as co-factors to TEAD transcription factors^22,23^. Several mutations in *YAP1* have been reported in humans as causing coloboma^24–28^. In *Yap1* homozygous mice, development was arrested at E8.5 due to requirement of YAP1 in yolk sac vasculogenesis and embryonic axis elongation^29^. *Yap1* heterozygote mice have complex ocular defects with microphthalmia, small palpebral fissures, anterior segment dysgenesis and cataracts^30^. Tissue specific ablation of *Yap1* in developing ocular tissues resulted in presumptive RPE acquiring anatomical and molecular characteristics of NR, including loss of pigmentation and pseudostratified epithelial morphology accompanied by ectopic induction of markers for NR progenitor cells^31^. Zebrafish *yap1* and *wwtr1* are specifically localized at 18 somite stage to the presumptive epidermis and notochord and play a role in the posterior body extension via regulating fibronectin^32^. Miesfeld et al., in 2015, have demonstrated via a transgenic line that *yap1* and *wwtr1* are expressed predominately in the developing RPE and lens, unlike homozygous knockout mice, zebrafish homozygous for *yap1* mutations survive, albeit with a variable phenotype depending on the allele. Although all reported alleles cause pigmentation defects in the eye, some alleles also cause coloboma. Double homozygous mutants for both *yap1* and *wwtr1* do not develop past 16 hours post fertilization (hpf). Embryos with one wildtype (WT) copy of *wwtr1* in a *yap1-/-* background survive with exacerbated pigment defects^32,33^.

In the current study, we seek to understand the mechanism by which *yap1-/-; wwtr1+/-*zebrafish develop coloboma. We demonstrate that *yap1* and *wwtr1* are expressed in the optic fissure, possibly in the pioneer cells, in addition to RPE, lens and NCC. *yap1* homozygote with one copy of *wwtr1* survive and have coloboma and pigmentation defects, more prominent at the optic fissure. RPE specific genes, though expressed in the RPE are patchy in nature, however, are totally excluded from the optic fissure area, same expression pattern is observed with the pigment genes which are dependent on the RPE specific genes. Pro-retinal genes, which are expressed in the bipotential optic cup are mis-expressed, which in turn affects the downstream retinal differentiation genes, resulting in transdifferentiation of retinal cell types in between the OF edges. This is the first observation in zebrafish where we see a transdifferentiation of NR at the expense of RPE. Our data suggests that in addition to a role in the specification of RPE, expression of *yap1* and *wwtr1* at the OF and in the RPE at the optic fissure edges is important for closure of the optic fissure. In the *yap1-/-; wwtr1+/-* mutant, due to the loss of these effectors the cells at the OF cannot attain “pioneer cell state”, where they can fuse, at the same time the cells cannot maintain their RPE state resulting in transdifferentiation of retinal cell types, causing a steric block and coloboma.

## Results

### yap1^-/-^; wwtr1^+/-^ mutants have coloboma and pigmentation defects

To understand the development of ocular defects in *yap1^-/-^; wwtr1^+/-^* mutants, we studied the expression of these two genes during ocular development. At 24 hpf *yap1* and *wwtr1* are expressed in the lens, OF, POM and faintly in the developing RPE (Supplementary Fig. 1A, A’, E, E’). Expression of these paralogues in the RPE cells become prominent by 36 hpf; importantly, their expression is also observed in the cells lining the OF—including, presumably, the pioneer cells, at the time of initiation of fusion (Supplementary Fig.1B, B’, F, F’). As the OF fusion proceeds the expression of *yap1* and *wwtr1* at the fissure edges recede, however its expression in the RPE is uniform at 48 hpf and lens expression becomes more circumscribed in the lens epithelium (Supplementary Fig. 1C, C’, G, G’). *yap1* and *wwtr1* expression ceases in the lens by 72 hpf but is still expressed in the RPE (Supplementary Fig. 1D, H).

**Figure 1:**
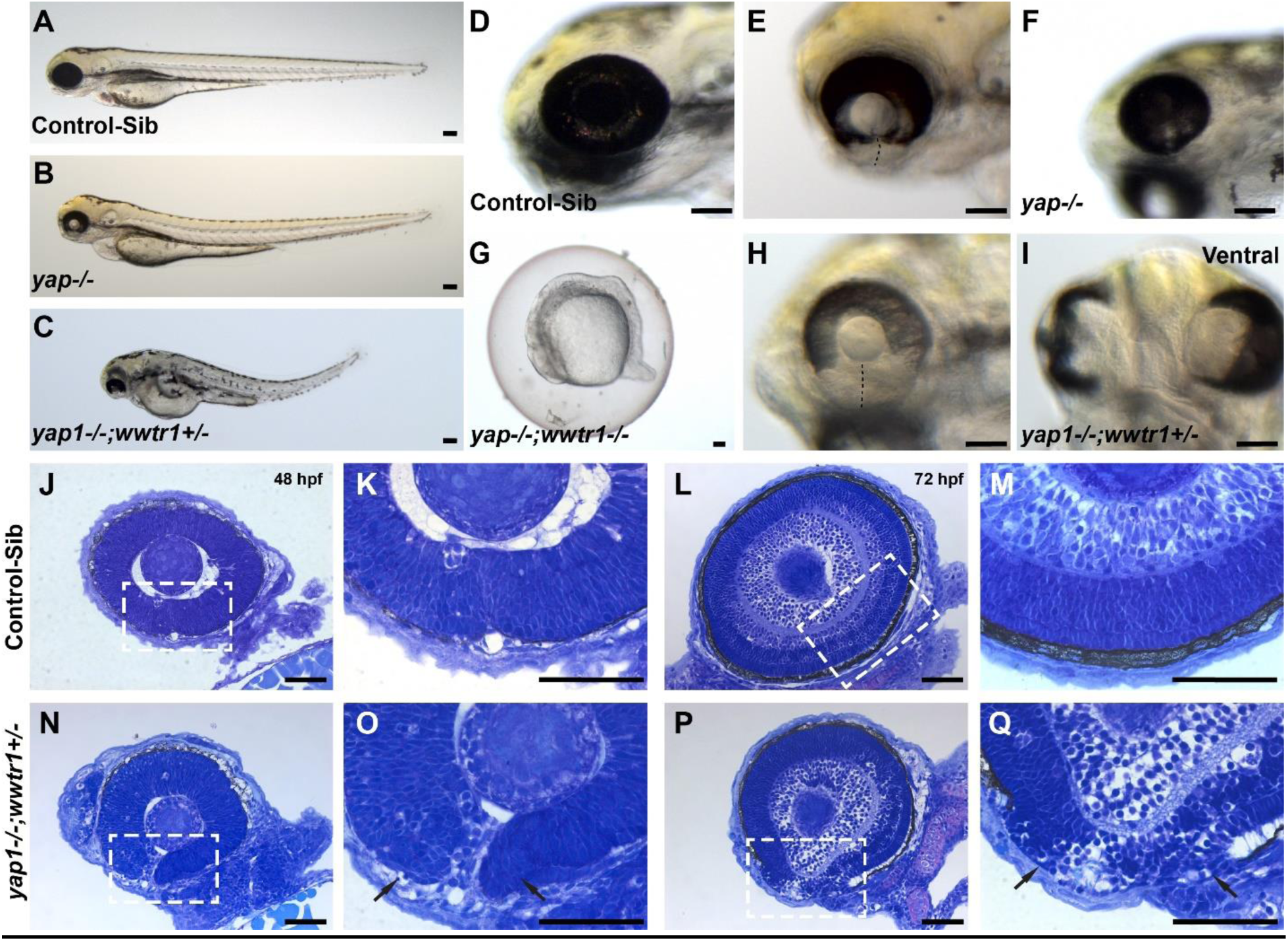
***yap-/-* and *wwtr1+/-* (YW) mutant embryos have coloboma and pigmentation defects:** At 72 hpf, eye morphogenesis and optic fissure (OF) closure is complete in control-sib embryos (A, D). *yap1^-/-^*embryos exhibit a spectrum of phenotypes including coloboma ± microphthalmia and pigmentation defects (B, E, F), which becomes more severe by heterozygous loss of *wwtr1* (*yap1^-/-^; wwtr1^+/-^* or YW mutants) (C, H, I). *yap1^-/-^; wwtr1^-/-^* embryos development arrests by 16 hpf (G). At 48 hpf, the edges of the OF are well-apposed in control-sib embryos (J, K), completely fusing by 72 hpf (L, M). The OF edges of YW mutants does not appose and presumed periocular mesenchyme cells can be seen between the edges at 48 hpf (N, O). OF edges do not fuse at 72 hpf, resulting in coloboma (P, Q). Reduced pigmentation in the presumptive retinal pigment epithelium (RPE) is noted in YW mutants at both time points (arrows in O and Q). Boxed areas in panels J, L, N and P are shown at higher magnification in K, M, O, and Q, respectively. Histological sections stained with toluene blue. All panels oriented with anterior to the left and are lateral views, I (ventral view). Scale bars 100µm (A-I) or 50µm (J-Q).

In our study, we used *yap1^bns^*^19^*^/bns^*^19^ and *wwtr1^bns^*^35^*^/bns^*^35^ (referred as *yap1^-/-^* and *wwtr1^-/-^*, respectively)^32,34^. *wwtr1^-/-^* mutants do not have any eye or pigmentation defects, appearing similar to control-sibs (Supplementary Fig. 2A-B’). However, at 72 hpf, *yap1^-/-^* embryos show variable loss of ocular pigmentation, with either bilateral coloboma and coloboma and/or microphthalmia, consistent with previous reports (Fig. 1A, B, D, E, Supplementary Table 1). Pigmentation defects were particularly notable ventrally, in the region of the OF. These phenotypes were augmented in the absence of one copy of *wwtr1* allele (*yap1^-/-^; wwtr1^+/-^*, referred from here as “YW mutants”) with notable reduction in ventral ocular pigmentation (Fig. 1C, H, I). Ocular abnormalities persist until at least five days in *yap1^-/-^* and YW mutants, suggesting that they are not due to developmental delay (data not shown). *yap1^-/-^; wwtr1^-/-^* embryos developmentally arrest by 16 hpf (Fig. 1G) and die by 26 hpf; these were not studied further. Histological analysis at 48 hpf revealed that control embryos have tightly apposed OF edges (Fig. 1J, K) compared to YW mutants where the OF edges are separated and we can observe presumed periocular mesenchyme (POM) cells (Fig. 1N, O). The OF of the control embryos are completely closed with a continuous layer of RPE and a well laminated retina at 72 hpf (Fig. 1L, M). In YW mutants, however, retinal lamination is disrupted and the OF remains open, resulting in coloboma (Fig. 1P, Q). Ventral pigmentation defects in the RPE are also notable on histology at both time points (Fig. 1O, Q, arrow).

### YW mutants do not have defects in optic cup morphogenesis

The first step in OF closure is the proper formation of the OC, we have studied OC morphogenesis in YW mutants via characterizing the expression patterns of known dorsal (*aldh1a2*, Fig. 2A, B), ventral (*vax2*, Fig. 2C, D), nasal (*foxG1a*, Fig. 2E, F) and temporal (*foxD1*, Fig. 2G, H) marker genes. No clear difference in the expression of these patterning genes was observed between control-sib and YW mutants at 24 hpf, suggesting that coloboma/microphthalmia in mutants is not caused by a major defect in ocular morphogenesis.

**Figure 2:**
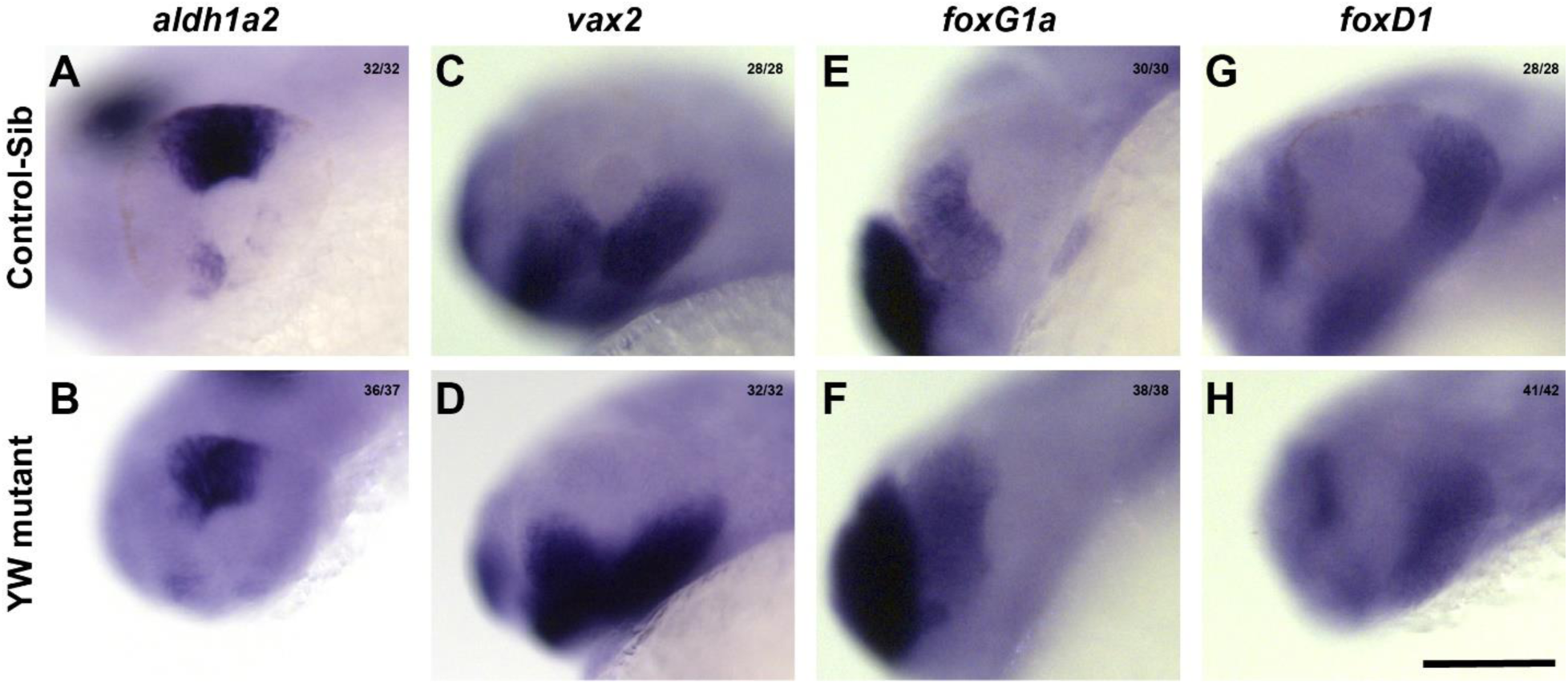
**Optic cup formation is not affected in YW mutants**: Expression of dorsal marker *aldh1a2* (A, B), ventral marker *vax2* (C, D), nasal marker *foxG1a* (E, F) and temporal marker *foxD1* (G, H) were similar between the control-sib and YW mutant embryos at 24 hpf. Scale bar is 100 µm.

### Coloboma in YW mutants is characterized by failure of OF edge apposition and loss of basement membrane breakdown

Another critical step in OF closure is the dissolution of the basement membrane (BM) between the apposed edges of the fissure, which is studied via laminin staining of the embryos. At 48 hpf, OF edges of the control embryos are well-apposed and the intervening BM dissolved in proximal and medial sections, however, BM can be consistently observed in the distal sections of the OC (Fig. 3A-C’). The OF edges of YW mutants are not completely apposed in any of the sections (Fig. 3D-F’). BM is completely breached and OF edges are fused in all the sections and the neuroepithelium appears continuous, consistent with fusion of control embryos at 72 hpf (Fig. 3G-I’) and coloboma in YW mutants, showing an intact BM with intervening tissue (Fig. 3J-L’, Supplementary Fig. 3). The tissue intervening between the OF edges shows an even more robust BM, with what appears to be extension from the temporal edge of the OC past the nasal edge (Fig. 3J, J’, K, K’). These observations raise the possibility that this intervening tissue may be providing a sort of steric hindrance to OF closure.

**Figure 3:**
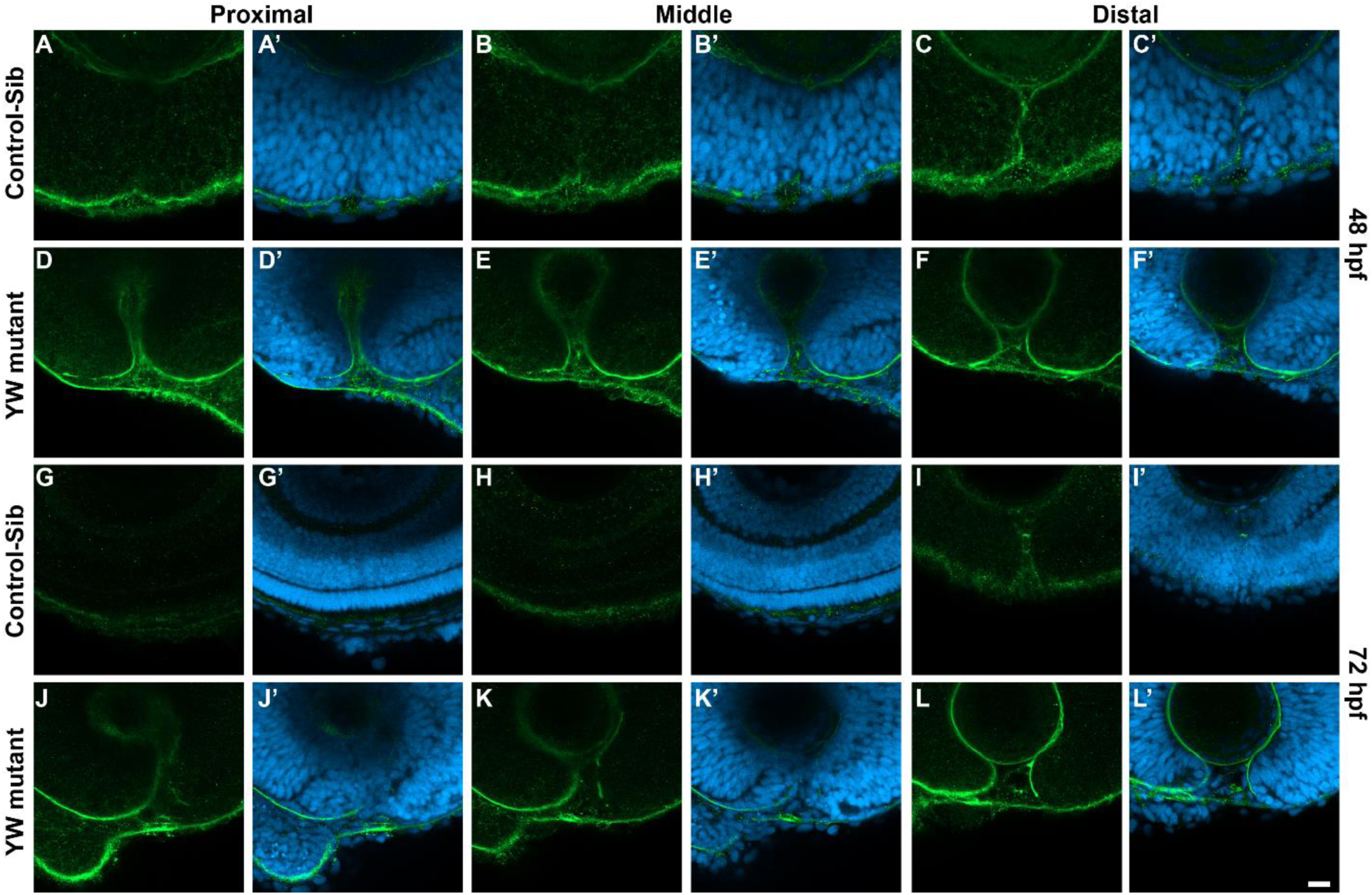
**Basement membrane does not breach in YW mutant embryos**: BM, as observed by laminin staining, is completely breached in proximal and middle sections (A-B’) and can only be seen in the distal sections at 48 hpf (C, C’) of the control-sib compared to YW mutants where we see laminin stain in all three sections (D-F’). At 72 hpf, laminin is absent in all the sections, indicating the fusion of OF edges in control-sib (G-I’) compared to YW mutants where we can see a very persistent laminin stain indicating coloboma (J-L’). Scale bar is 20 µm.

To better study this phenotype, we studied the expression of *ntn1a* which marks the OF edges^35,36^ via in situ analysis. *ntn1a* has prominent expression at the OF edges in the control-sib at 24 hpf (Fig. 4A); as OF fusion proceeds through 36 and 48 hpf, its expression recedes (Fig. 4B, C). By contrast, expression of *ntn1a* in YW mutants is mis-expressed at 24 hpf with more diffuse expression in the ventral OC, often with notable extension along its ventro-temporal edge (Fig. 4D). Faint expression is observed at the OF edges at 36 hpf and by 48 hpf a clear separation of the OF edges with intervening tissue is delineated by separation of the *ntn1a* signal, again suggesting a tissue block/cell proliferation between the OF edges (Fig. 4E, F).

**Figure 4:**
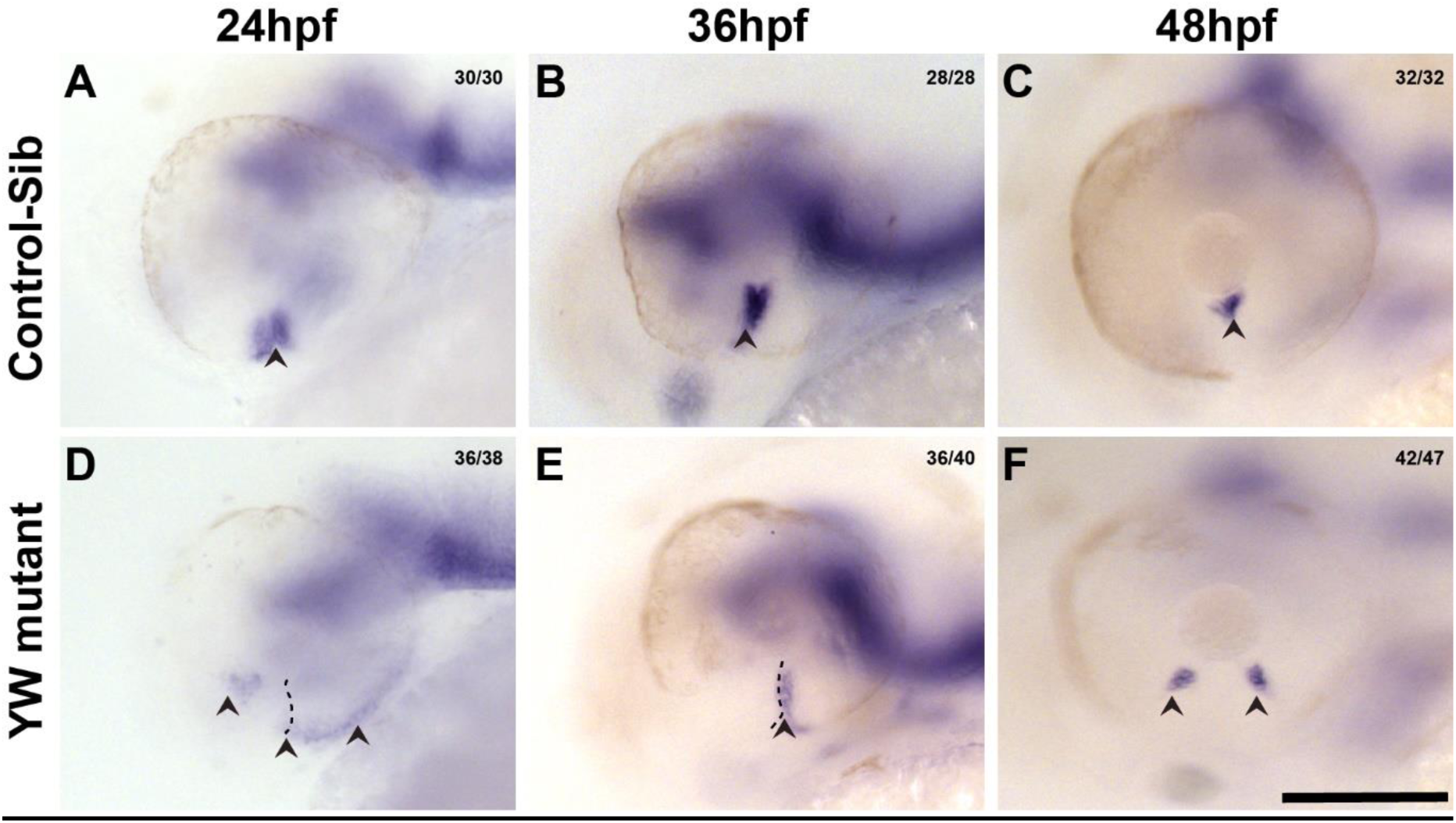
***<u>ntn1a</u>*<u>, which marks the OF edges is mis-expressed in YW mutant embryos</u>:** In control embryos, *ntn1a* is expressed in the OF edges at 24 hpf and 36 hpf (A, B). As fusion process proceeds, its expression recedes to a small patch in the inner aspect of the optic cup by 48 hpf (C). YW mutant embryos *ntn1a* expression is seen at the sides of the OC (D) rather than OF edges at 24 hpf, and by 36 hpf expression is seen at the OF edges (E). However, by 48 hpf, we see a tissue in between the expression of *ntn1a* (F). Arrow heads point to the *ntn1a* expression. Scale bar is 100 μm.

### The tissue between the OF edges is not due to overgrown optic stalk

We next sought to characterize the nature of the cells present in the OF. *George et al.* have recently reported that the coloboma in the *rerea* mutant zebrafish is caused by a shh-dependent overgrowth of the OS neuroepithelium. *fgf8* is a known morphogen to pattern the NR and the OS and to promote retinal ganglion cell differentiation in zebrafish and chick^37^*. fgf8 is* expressed in the forebrain in addition to the OS at 24 hpf and 36 hpf in the control-sib. Expression is diminished in the OS in the control-sib embryos, with the differentiation of retinal ganglion cells and the formation of the optic nerve at 48 hpf (Supplementary Fig. 4A-C). YW mutants have similar expression of *fgf8* in the brain, however, expression in the OS is sparse at 24, 36 and 48 hpf (Supplementary Fig. 4D-F). We observe the same pattern of expression with the *pax2a* gene which is under the control of *fgf8* and important for OF fusion (data not shown). Sonic hedgehog signaling which governs the OS formation has normal expression in the YW mutants compared to the control-sib (Supplementary Fig. 5A-B’). These observations confirmed that coloboma in YW mutants is not due to an overgrown OS.

### Expression of RPE and pigment-related genes are reduced at the OF in the absence of yap1 and wwtr1

*yap1* and *wwtr1* have well established roles in RPE biogenesis in the developing OC of zebrafish. A subset of presumed RPE cells; the “pioneer cells”^9^ are the first to make contact along the OF edges and initiate fusion. RPE development in YW mutants is studied via in situ analysis of the well-characterized RPE specific genes, *mitfA* and *tfec*. In control embryos at 36 and 48 hpf, *mitfA* is expressed in the RPE on the outer edge of the OC and NCCs (Fig. 5A, C); *tfec* shows similar RPE and NCC expression at both time points studied, in addition, it is also expressed in the OF. (Fig. 5E, G). In contrast, YW mutants have patchy expression of both genes, albeit in a similar pattern to controls. We note that expression in the OF area is specifically missing for both genes at both time points (Fig. 5B, D, F, H). Similarly, *otx2a,* another transcription factor which plays an important role in the formation of RPE, is widely expressed throughout the OC at 36 and 48 hpf in control embryos (Supplementary Fig. 6A, B). YW mutants show a similar but sometimes patchy pattern expression in OC, albeit with absent expression in the OF region at both time points (Supplementary Fig. 6C, D, E). RPE specific genes, especially *mitf*, play a role in promoting melanocyte and RPE differentiation via regulating transcriptional control of pigmentation genes. *dct* a member of tyrosinase gene family was expressed all over the eye at 24 hpf and 36 hpf, consistent with normal pigmentation in control-sib embryos (Fig. 5I, K) compared to YW mutants that exhibit sparse and reduced expression around the fissure edges at 24 hpf (Fig. 5J). By 36 hpf a clear loss of *dct* expression at the OF edges is noted (Fig. 5L). These observations are consistent with the pigmentation defects observed in the eye, especially at the OF edges, indicating that YW mutants have pigment defects particularly pronounced at the OF edges.

**Figure 5:**
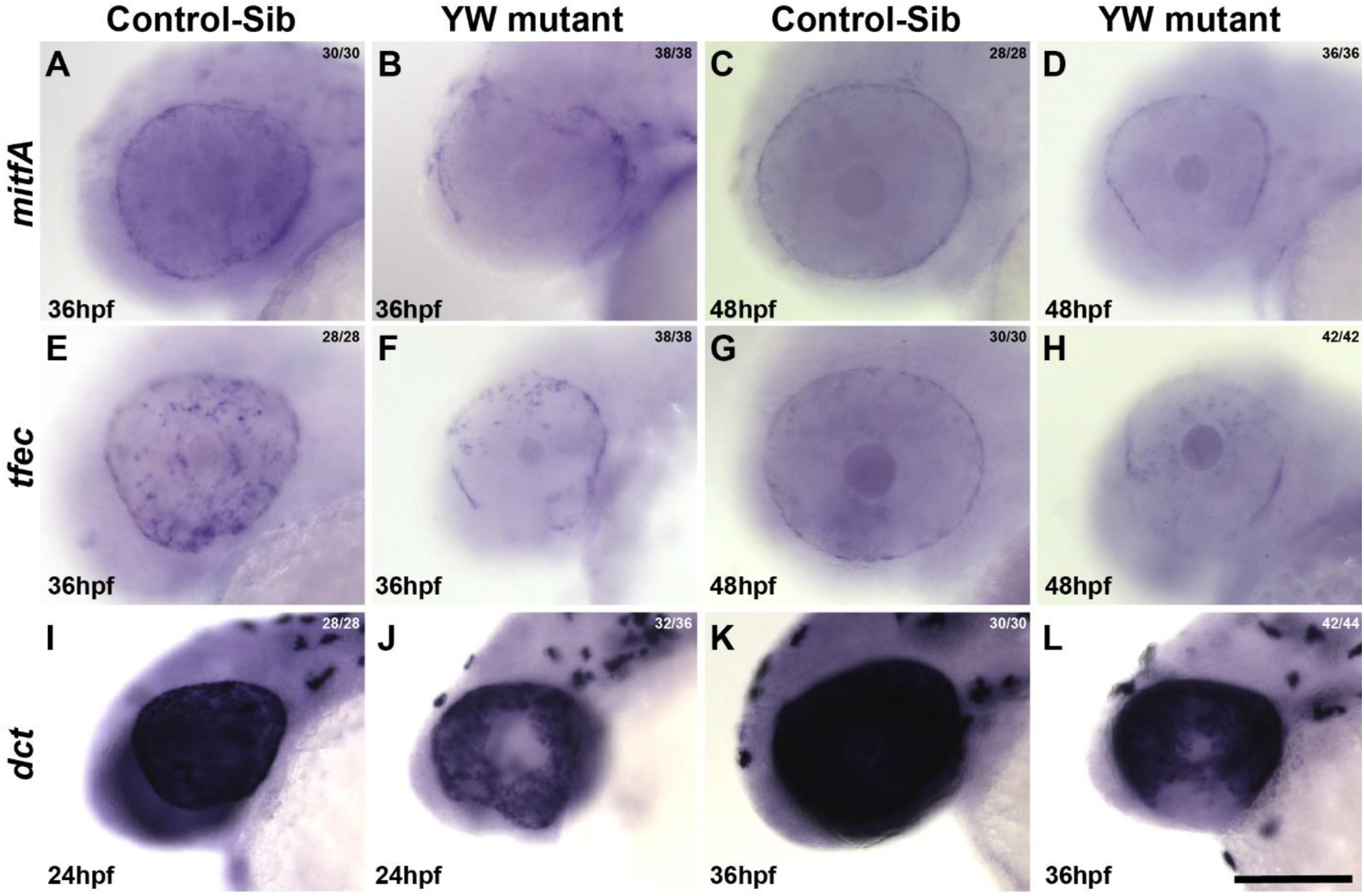
RPE and melanogenic specific gene expression is specifically excluded from the OF area of YW mutant embryos: *mitfA* is expressed in the NCC and RPE at 36 hpf and 48 hpf (A, C) in control-sib embryos. In YW, expression of *mitfa* is patchy in the RPE, NCC and is undetectable at the OF edges at both 36 hpf and 48 hpf (B, D). *tfec* is expressed in the OF edges, NCC and RPE at 36 hpf (E), RPE and NCC at 48 hpf (G) in control-sib embryos. Like *mitfA*, RPE expression is patchy and is undetectable at the OF edges (F, H) at both 36 and 48 hpf in YW embryos. *dct* is expressed broadly in the OC at 24 and 36 hpf in control embryos (I, K). Its expression, however, is reduced at the OF edges in YW embryos at 24 hpf (J). By 36 hpf its expression is undetectable at the edges of the OF (L). Scale bar is 100 µm.

### Coloboma is accompanied by the presence of differentiated NR, rather than RPE tissue at the edges of OF in YW mutants

We next evaluated the possibility of presumptive NR as the source of cells remaining in the OF. The transcription factor *pax6* is expressed at 24 hpf throughout the optic cup, in both pNR and pRPE (Fig. 6A, B); YW mutants show a similar pattern of expression, albeit with a relative increase in expression (Fig. 6G, H). During the later stages of OC development (36 hpf), *pax6* expression is lost in RPE (Supplementary Fig. 7A) and later becomes restricted to ganglion and amacrine cell layers of the developing retina of control embryos (Supplementary Fig. 7C, E). Expression at 36 hpf in the YW mutants is throughout the OC, except for reduced expression in the OF and along with ventro-nasal and temporal regions of the OC (Supplementary Fig. 7B, arrows and Asterix). At 48 hpf, *pax6* expression becomes restricted to the ganglion and amacrine cells of the retina but is specifically excluded in the optic fissure region (Supplementary Fig. 7D, Asterix). Surprisingly by 72 hpf YW mutants regain the expression of *pax6* at the OF, which extends into the fissure edges. (Supplementary Fig. 7F, arrowheads). *vsx2,* an early pNR marker, is normally expressed at 36 hpf throughout the OC and becomes restricted to the Müller glia and bipolar cells by 48 hpf in the control-sib (Fig. 6C-F). The expression of *vsx2* in the optic cup of YW mutants is similar in pattern to the control-sib at 36 hpf, however they have a sparse or patchy expression at the OF edges compared to the controls (Fig. 6J, Asterix). By 48 hpf there is no expression of *vsx2* at the fissure edges (Fig. 6I-L). Since *vsx2* plays an important role in the development of other retinal cell types, we furthered our study into other cell types. Ganglion cells are the first retinal cell type to develop and can be seen at 36 hpf adjacent to the lens, as observed by *alcama* in situ stain. As OC development progress, they can be seen as encircling the lens by 48 hpf (Fig. 6M, N). YW mutants display a similar pattern, but also extend into the OF at 36 hpf, becoming more prominent by 48 hpf (Fig. 6S, T). The amacrine and red/green photoreceptor markers Huc/D and Zpr1 are expressed circumferentially at 72 hpf in the inner and outer retina, respectively (Fig. 6O-R). However, in YW mutants these markers extend into the OF margins (Fig 6.U-X, arrowhead). Moreover, we have observed that the photoreceptors do not develop well in the regions which do not have a well-developed RPE, disrupting retinal lamination (Supplementary Fig. 8A-B’).

**Figure 6:**
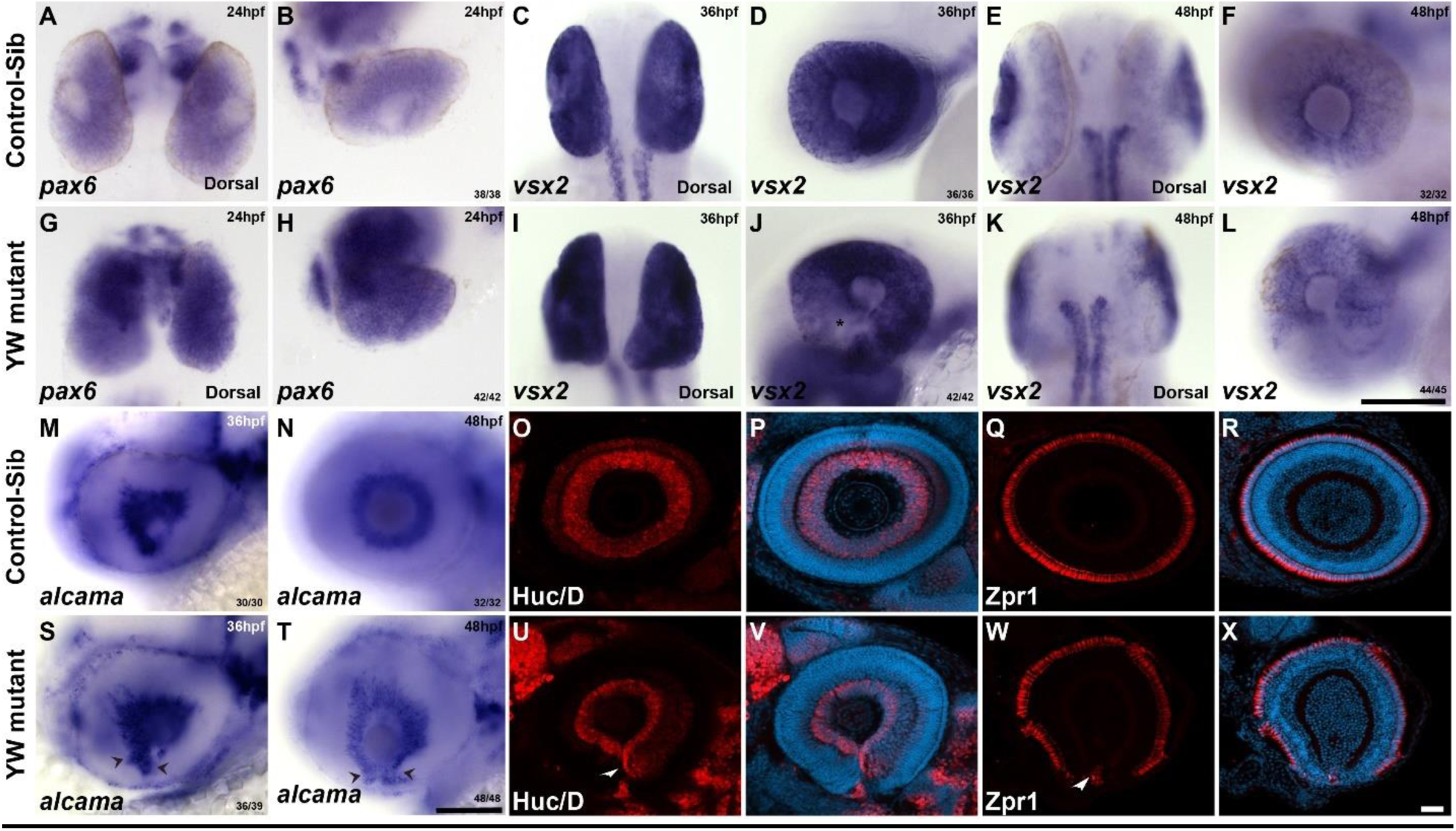
Early retinal markers are upregulated, and late retinal markers are expressed between the OF edges: In control-sib embryos, *pax6*, which is expressed in the bipotential cells of the OC, is expressed throughout the OC at 24 hpf (A, B). This expression is notably increased in YW mutant embryos (G, H). *vsx2* is expressed broadly in the presumptive neural retina of the OC at 36 hpf (C, D) and becomes restricted to Müller glia and bipolar cells in the retina by 48 hpf (E, F). However, YW embryos have a patchy expression in the nasal side of the OC at 36 hpf (I, J, Asterix); by 48 hpf its expression is though restricted to bipolar cells and Müller glial cells, its expression is notably excluded from the OF edges (K, L). *alcama* is expressed in the ganglion cells at 36 and 48 hpf in control-sib embryos (M, N). In YW embryos its expression extends ventrally into the OF edges (S, T, arrow heads). In control embryos at 72 hpf, the amacrine cell marker Huc/D (O, P) and the photoreceptor marker Zpr1 (Q, R) are expressed circumferentially in the inner and outer retina, respectively. In YW embryos both Huc/D and Zpr1 are expressed in a similar pattern but extend ventrally between the OF edges (U-X, arrow heads). Scale bars are 100 µm for panels A-T and 20 µm for panels O-X. The embryos are dorsal where mentioned, rest of the embryos were oriented laterally, anterior towards left.

## Discussion

In this study, we sought to understand, specifically, the role of the Hippo signaling effectors *yap1*/*wwtr1* in optic fissure closure and the pathogenesis of coloboma. We demonstrate that *yap1* and *wwtr1* are dynamically expressed during zebrafish eye development—including at the edges of the optic fissure—and that the coloboma caused by loss of *yap1* function is exacerbated by mutation of one *wwtr1* allele. This phenotype occurs despite normal optic cup morphogenesis, with clear separation of the basement membranes of the OF edges. Rather, we note misexpression of OF markers such as *ntn1a*, reduction of RPE-associated markers (*mitfA, tfec*), and abnormal expression of early retinal markers (*pax6, vsx2*) at the closing edges of the OF in YW mutants—all of which likely put presumed pioneer cells in a developmental state inconsistent with fusion. This pattern of gene expression also likely explains the overgrowth of non-pigmented, poorly laminated NR tissue at the fissure which itself may sterically interfere with fusion.

Previous reports have focused on the role of *wwtr1* and *yap1* in early ocular development, specifically in RPE determination^33^. We extend these findings, by further exploring the mechanism of coloboma. We, as others, noted that loss of *wwtr1* does not result in an ocular phenotype, but loss-of-function of one *wwtr1* allele exacerbates the coloboma and pigmentation defects of *yap1^-/-^* embryos^32,33^. Different *yap1* alleles have been shown to cause different phenotypes in zebrafish. For example, the *yap^n1^*^13^ mutation alters *yap1* splicing, creating a truncated protein, leading to coloboma without significant pigment abnormalities. By contrast, the *yap^mw48^* allele contains a frameshift allele that results in loss of *yap1* immunoreactivity (i.e., presumed loss-of-function) and exhibits pigment defects but lacks coloboma^33,38^. The *yap1* homozygotes used in our study *yap1^bns19^* harbor an early frameshift mutation in the TEAD-binding domain^32^ and have both coloboma and pigmentation defects, consistent with human mutations of *YAP1*^25,27,28,39^.

Morphogenetic coloboma presents itself as a gap between the optic fissure edges due to defective optic cup morphogenesis, owing to the failure of genes responsible for earlier stages of OC formation^9^. Coloboma observed in YW mutants is not morphogenetic in nature, as the optic cup forms relatively normal and there is no clear disruption in classic developmental patterning genes. We and others have also found that mutation of genes such as *ptch1* and *rerea* can result in coloboma, from a widened optic stalk leading to abnormal optic cup morphology; this phenotype was not seen in YW mutants, consistent with reduced *pax2a* and *fgf8* expression^40–42^

The abnormal pattern of ventral RPE development suggests, rather, that the mechanism of coloboma is due to a failure of the cells at the edges of the optic fissure to attain a developmental state conducive to fusion. Eckert et al., have noted that the first cells of the closing edges of the OF to make contact are “pioneer cells” that are morphologically similar to presumptive RPE cells and that do not express the neural retina marker *rx2*^9^. The decreased ventral expression patterns we observe in RPE markers *mitfa*, *tfec* and *otx2* coupled with increased expression of the early NR marker *pax6* suggested that the region containing these pioneer cells fails to attain this pattern. Indeed, these ventral cells go on to form poorly laminated NR-like tissue that expresses markers for multiple NR cell types that “overgrows” across the OF with time, we cannot exclude the possibility that physical disruption of OF apposition also contributes to coloboma.

This pattern of gene expression leading to formation of NR at the expense of RPE has been noted in multiple mouse models, however this is the first observation in zebrafish. For example, mutations in the RPE specific isoforms of *Mitf* gene in mice results in microphthalmia and coloboma, with RPE transdifferentiating into NR and retinal degeneration^43,44^. Similarly, *Otx2^+/-^; Otx1^-/-^* mouse embryos develop NR tissue at the expense of RPE^45,46^. In zebrafish neither *mitfa*^47,48^ nor *tfec* mutations result in coloboma but do have pigmentation defects. *tfec* mutant eyes are mildly microphthalmic, with normal optic cup development^49^ and *mitf/tfec* double mutants have coloboma^49^. *otx2* and its orthologs, *otx1a* have a role in RPE specification in zebrafish and knock down of both results in coloboma and ventral pigmentation defects, but not with abnormal NR formation, as seen in mouse mutants^50^.

*pax6* regulates *vsx2* and, together, these commit bipotential cells in the OC to a NR fate^51^, with *vsx2* and *mitf/tfec* mutually inhibiting each other^52^. Surprisingly, we do not observe an upregulation of *vsx2* at the OF edges where *mitf/tfec* expression is reduced. This may be explained by the transient downregulation of *pax6* we observe at 36 hpf and 48 hpf in the OF with deficient hippo signaling as the major cause of reduced *mitf/tfec* expression.

The tissue that grows across the OF differentiates into some, but not all NR cell types. Specifically, we note this tissue expresses ganglion, amacrine and R/G photoreceptor cell markers, but not the bipolar cell marker *vsx2*. We posit that the expression of ganglion, amacrine and photoreceptor markers is made possible by the subsequent rebound in *pax6* expression by 72 hpf^53^. Presumably the abnormal lamination we observe is also due to this “imbalance” of NR cell types in the ventral eye.

Lastly, we note that Hippo signaling is a well-recognized regulator of cell movement as well as of RPE development^38,54^. The images in this study are static and, as such, we cannot assess whether at least some of the abnormal cell fates we observe are secondary to disruption of the normal movement of cells from the forebrain (extended evagination) which are required for optic stalk and optic fissure formation^55^. The diffuse expression of the OF marker *ntn1a* we observe at 24 hpf--with later concentration near the OF edges—may be consistent with this mechanism. Normally, cells in outer layer of the OC migrate around its rim to integrate into the inner retinal layers^3–5^. We speculate that abnormal hippo signaling may compromise such movements, effectively “trapping” retinal cells at the OF edges and creating a potential physical block to closure. An interesting follow-up study to this work would therefore be to image YW mutants using “4-D live imaging” that can quantitate cell movements.

## Material and Methods

### Fish maintenance and zebrafish strains

*Danio rerio* were maintained under standard conditions. Embryos were staged according to Kimmel et al., 1995^56^. ABTL stocks, *yap1^bns19^* and *wwtr1^bns^*^35^ ^32,34^ were used in the study and are referred to as *yap1^-/-^* and *wwtr1^-/-^*, respectively, in this paper (Supplementary Table 2). Experiments were carried out in accordance with National Eye Institute, Animal Care and Use Committee Protocol Number NEI-648.

### Plastic Sectioning and methylene blue staining

Plastic sectioning is carried out according to following the protocol^57^. Plastic sections were stained in 1% Toluidine blue in 1% borate.

### Zebrafish in situ hybridization

Embryos were fixed in 4% paraformaldehyde (PFA) overnight at 4°C and dehydrated in methanol for 1h at −30°C. The embryos were rehydrated, treated with proteinase-K and re-fixed with 4% PFA. Pre-hybridization and hybridization were carried out at 65°C. Primers were designed with T7 promoter in the reverse primer (Supplementary Table 3) and the genes were amplified, which were used as templates to make DIG labeled RNA probes. These probes were synthesized using a DIG labeling kit (Millipore-Sigma, 112770739) following manufacturer’s protocol. Samples were hybridized overnight with RNA probes at 65°C, washed, incubated with Anti-DIG antibody (Millipore-Sigma, 1109327490); color was developed using BCIP/NBT substrate (Millipore-Sigma, 11681451001) in alkaline phosphatase buffer. Embryos were imaged with Leica DM6 dissecting microscope.

### Immunohistochemistry

Embryos were fixed in 4% paraformaldehyde overnight at 4°C. After washing with 1× PBST, the embryos were blocked in blocking buffer (2% donkey serum, 0.1% Tween and 1× PBS). Primary antibodies used were at 1:100 dilution in the blocking buffer, Laminin (Sigma-L9393), Huc/D (Invitrogen, A21272), Zpr1 (ZIRC) (Supplementary Table 4). Primary incubations were carried out overnight at 4°C. After the primary incubations the embryos are washed several times with PBST and stained with nuclear stain Hoechst 33342 (Invitrogen, 2306347) and Alexa Flour^TM^ 488 (Invitrogen, A21206) for two hours at room temperature. Washed several times in dark at room temperature and transferred to glycerol for storing and imaging. Zpr1 stain was carried out on cryosections and Huc/D and Laminin on whole mount embryos, which were imaged on Zeiss LSM 800.

## Supporting information

Supplemental Tables for Yap1 wwtr1 article

## Acknowledgments

This study was supported by the Intramural Research Program of the National Institutes of Health. We thank Prof. David Kimmelman, University of Washington, for sharing the *yap1^bns19^* and *wwtr1^bns35^* zebrafish mutants. We thank the Aquatic facility staff of NEI, for their help and support.

## Author Contribution

UMN and BPB designed the study, experimental design, drafting of the manuscript and overall project support. UMN performed experiments and analyzed the data. DSM performed phenotype analysis, SS and RA helped with genotyping of the mutants.

---

**Supplementary Fig. 1:**
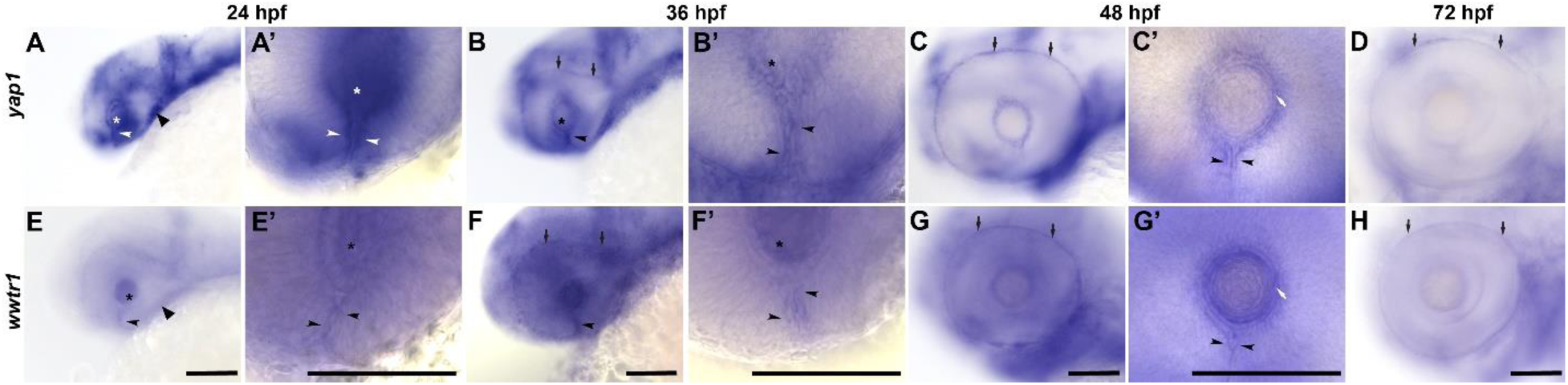
**Expression of *yap1* and *wwtr1* in zebrafish eye at various developmental stages**: *yap1* and *wwtr1* are expressed in the OF (arrowhead), lens (Asterix) and periocular mesenchyme cells (closed arrowhead) at 24 hpf (A, A’, E, E’). By 36 hpf (B, B’, F, F’) expression is seen in the RPE (arrows) in addition to OF (possibly in pioneer cells, arrowheads) and lens. At 48 hpf, expression at the OF recedes close to lens; becomes prominent in the RPE (C, D), and the expression in lens becomes more restricted to lens epithelium (white arrows in C’ and G’). By 72 hpf its expression in the lens epithelium is lost and is only present in the RPE (D, H). A’, B’, C’, E’, F’, G’ are higher magnification images of the optic fissure area of A, B, C, E, F, G respectively. Scale bar is 100 μm. Anterior towards left.

**Supplementary Fig. 2:**
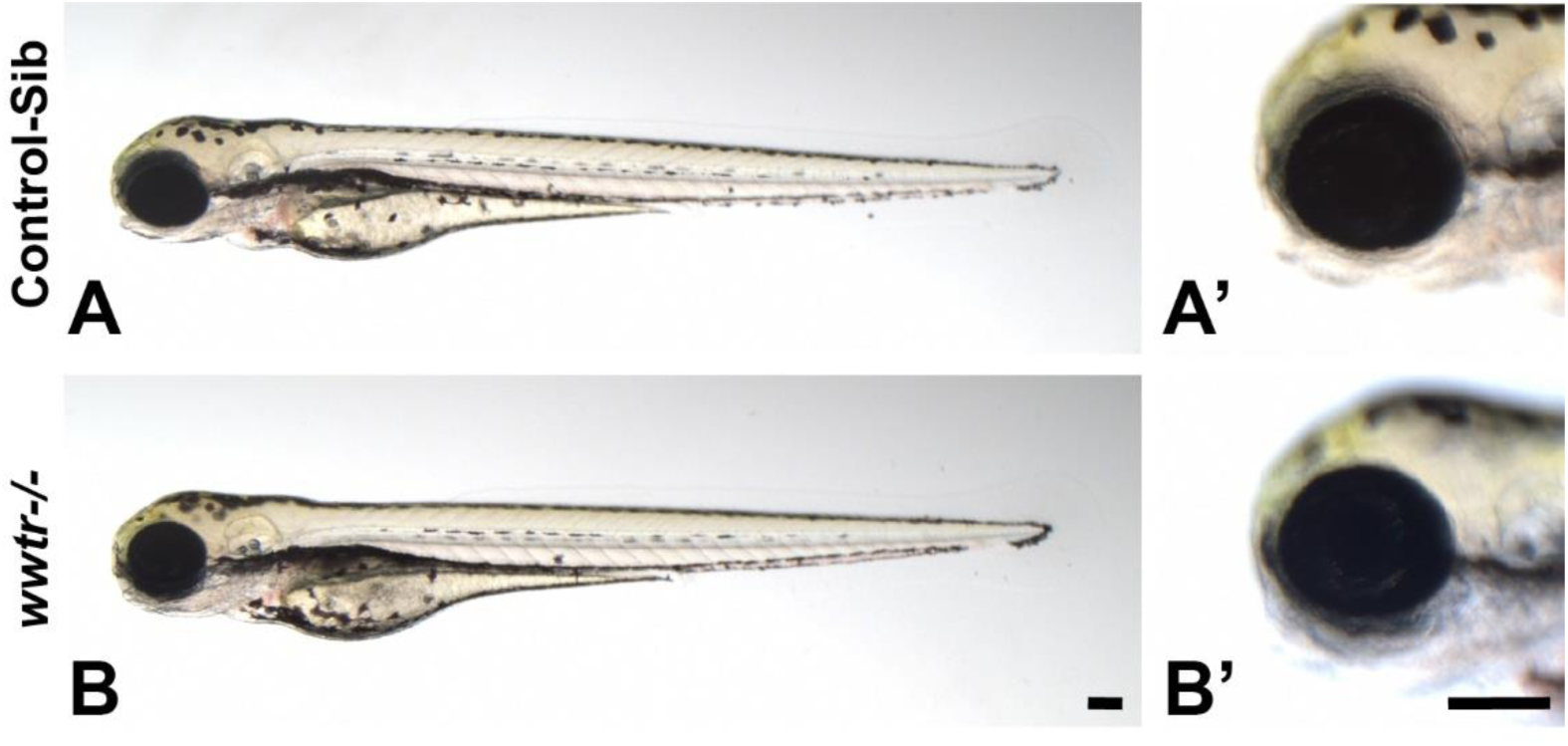
*wwtr1-/-* mutants do not have ocular or pigmentation defects: Phenotypic difference was not observed between *wwtr1-/-* (B, B’) compared to control-sib at 72 hpf (A, A’). Scale bar is 100 μm. Phenotype images were taken at 72 hpf.

**Supplementary Fig. 3:**
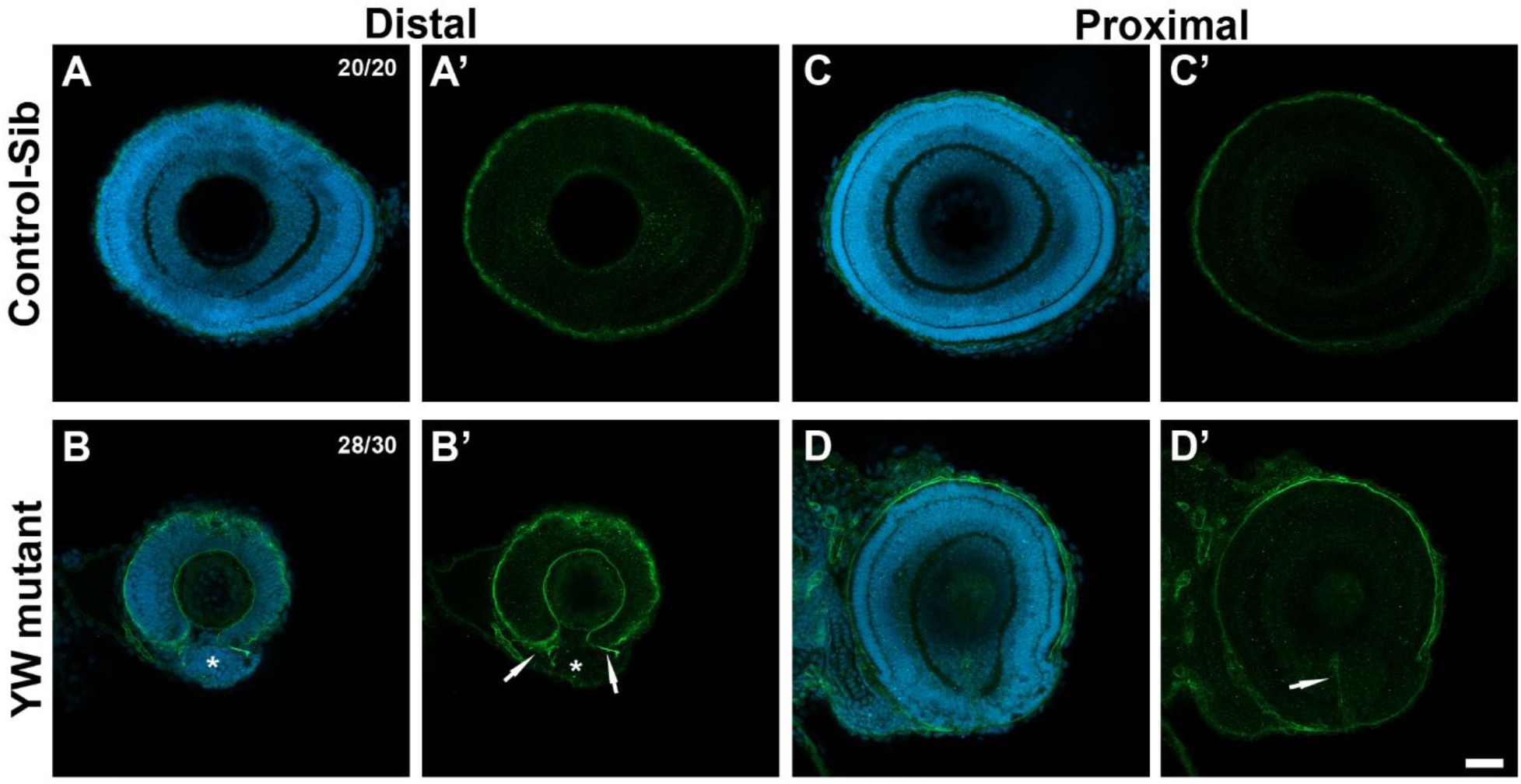
Failure of OF closure in YW mutant embryos is due to a possible tissue hinderance: Basement membrane is breached as indicated by the absence of laminin staining in the distal and proximal sections and OF is fused in the control-sib at 72 hpf (A-C’). However, laminin staining in YW mutants delineate clearly visible optic fissure edges (arrows) with a tissue between the edges (Asterisk) in the distal sections (B- D’). Scale bar is 20 µm.

**Supplementary Fig. 4:**
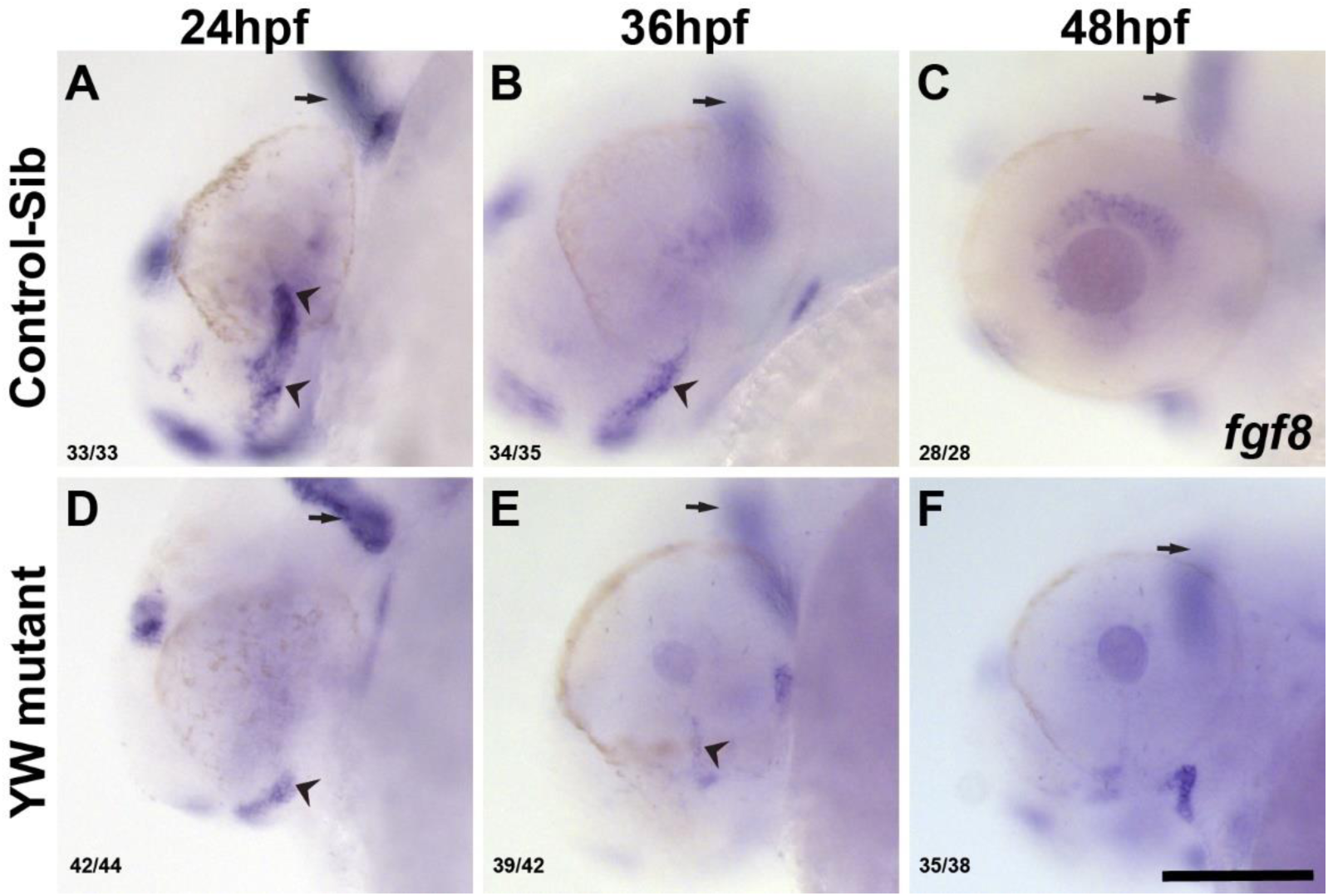
Defects in OS formation in YW mutants: Control-sibs have expression of *fgf8* in the OS at 24 hpf (A) and 36 hpf (B); by 48 hpf its expression in the OS is reduced and can be seen in the retina (C). YW mutant embryos, by contrast, demonstrated reduced expression of *fgf8* at 24 hpf (D), minimal expression levels at 36 hpf (E) and faint expression at the OS and retina at 48 hpf (F). Arrows and arrow heads point to the expression of *fgf8* expression in the brain and OF respectively. Scale bar is 100 µm.

**Supplementary Fig. 5:**
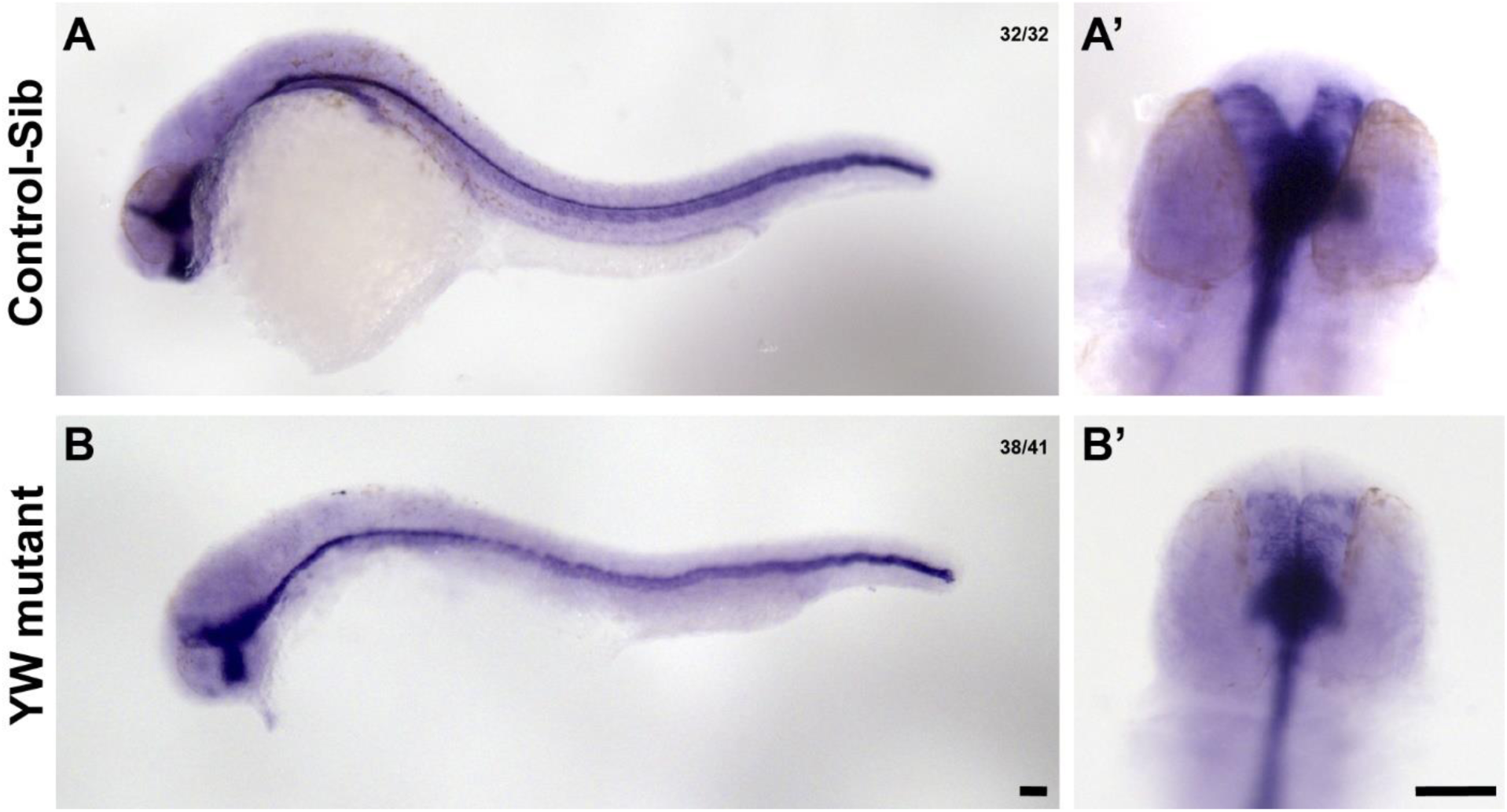
**Midline expression of *shha* is not affected in the YW mutants**: Midline expression of *shha* in the control-sib is similar to the YW mutants at 24 hpf (A-B’). Scale bar is 100 µm.

**Supplementary Fig. 6:**
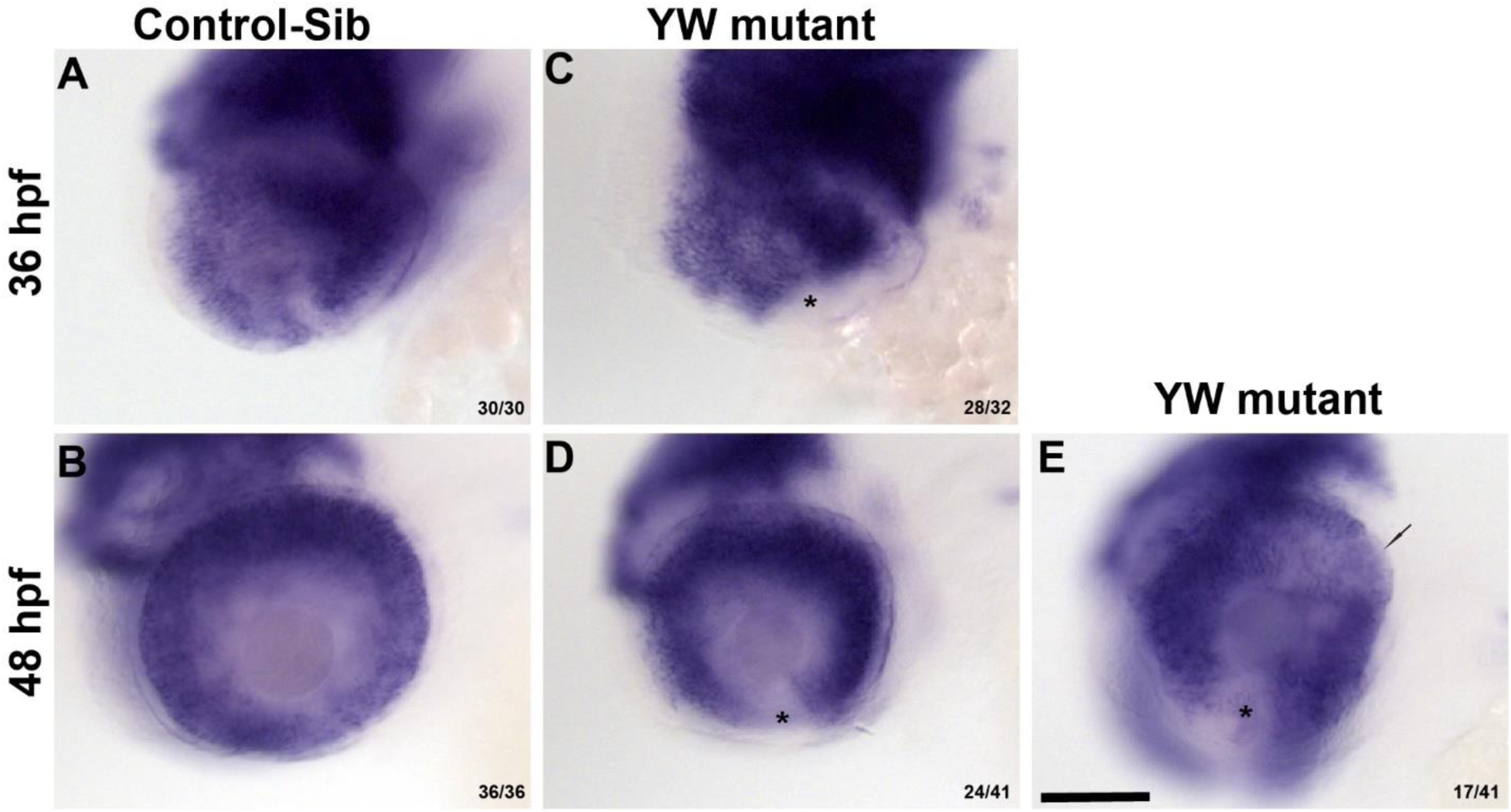
**RPE specific Transcription factor *otx2a* is not expressed at the OF in YW mutant embryos**: *otx2a* is expressed throughout the OC in 36 hpf and 48hp in control-sib embryos (A, B). YW mutants at 36 hpf exhibit either: 1) broad *otx2a* expression with reduced expression at the OF edges (C), which becomes evident as a gap by 48 hpf (D, Asterix) or 2) patchy expression of *otx2a* in the OC with reduced OF expression (E, arrow). Scale bar is 100 µm.

**Supplementary Fig. 7:**
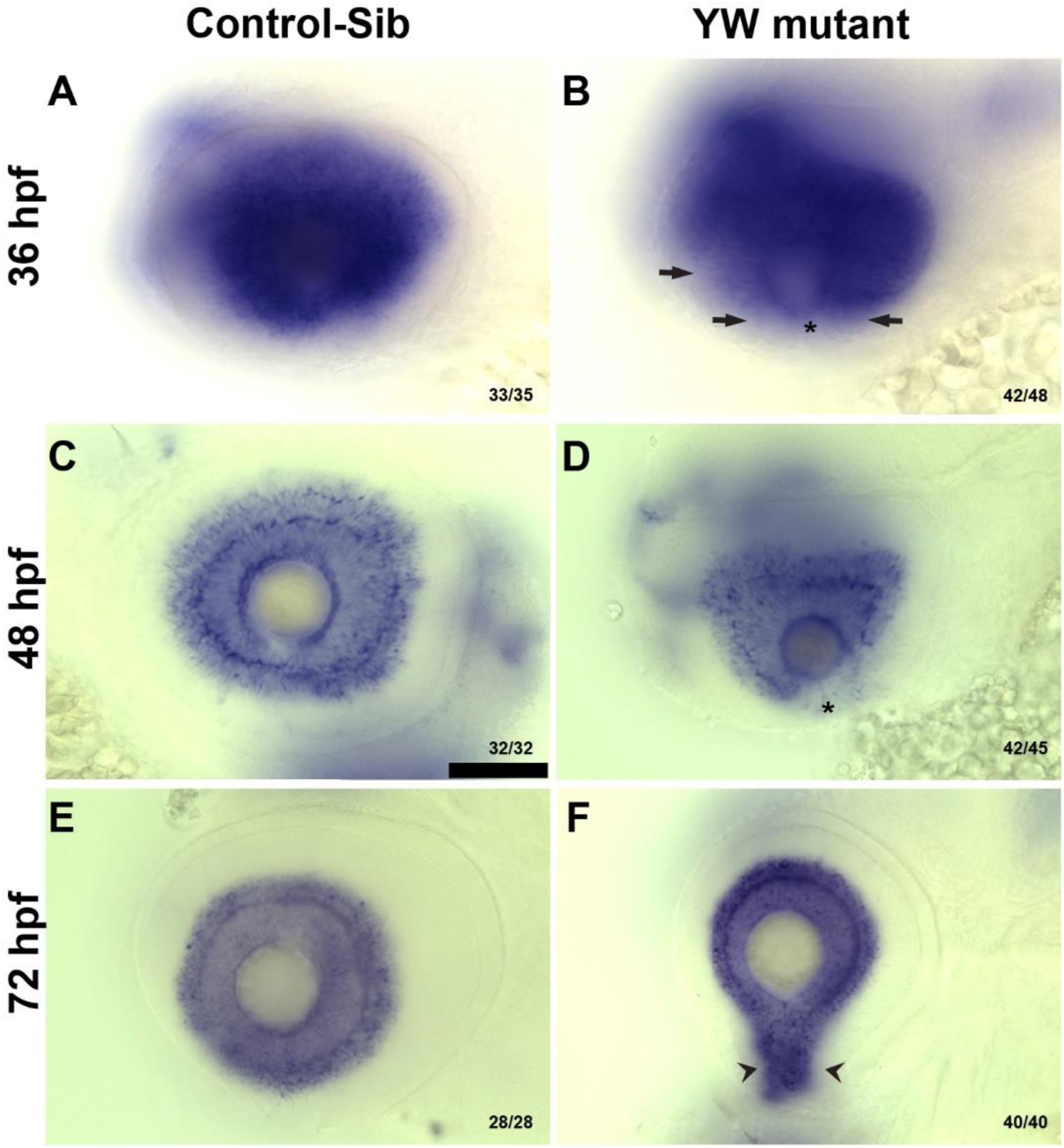
*pax6,* an early marker of the bipotential OC is mis-expressed in the YW mutants: *pax6* is expressed in the ganglion and amacrine cells and their processes at 36 hpf (A), 48 hpf (C) and 72 hpf (E) in the control-sib. YW mutants, have *pax6* expression all over the OC, no expression is observed at the OF and patchy expression on the sides on the OC at 36 hpf (B, Asterix, arrows). Expression of *pax6* is not observed at OF edges and the temporal lobe of the OC have patchy expression at 48 hpf (D, Asterix) and by 72 hpf, the mutants regain expression which extends into the open OF (F, arrowheads). Scale bar is 100 µm.

**Supplementary Fig. 8:**
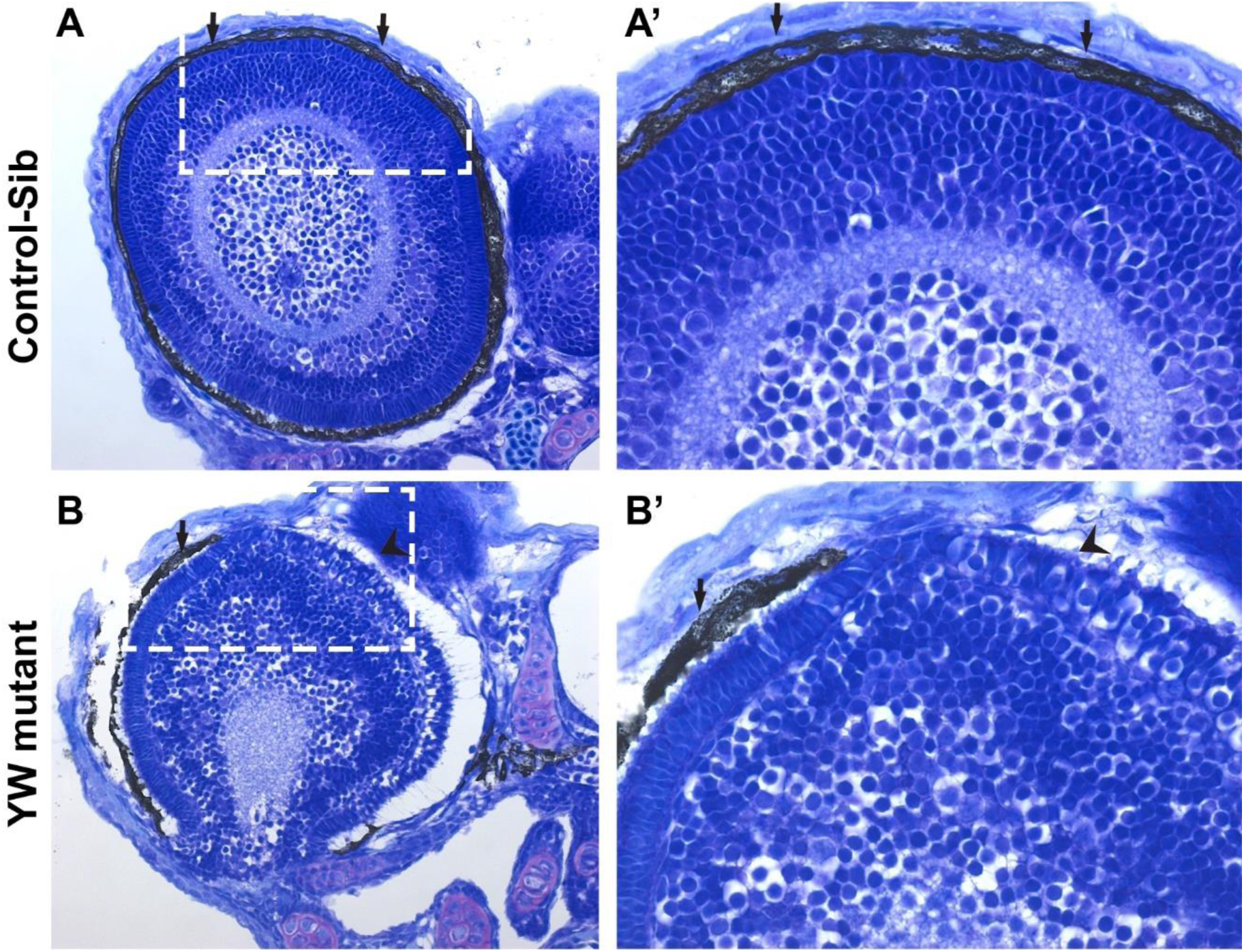
Patchy RPE defects in YW mutant embryos is accompanied by abnormal NR lamination: Histology of the OC at 72 hpf show a well formed RPE and NR in a control-sib (A, A’). However, in YW mutant embryos, retinal lamination is affected and RPE formation is patchy. In the regions of OC having pigment defects, photoreceptor formation is affected (B, B’). A’ and B’ are high magnification of boxed area in the A and B respectively. Arrows point to areas of retina having RPE and arrowhead to the area without RPE. Scale bar is 50 µm.

## References

1 Bibliowicz, J., Tittle, R. K. & Gross, J. M. Toward a better understanding of human eye disease insights from the zebrafish, Danio rerio. Prog Mol Biol Transl Sci 100, 287–330 (2011). 10.1016/B978-0-12-384878-9.00007-8

2 Richardson, R., Tracey-White, D., Webster, A. & Moosajee, M. The zebrafish eye-a paradigm for investigating human ocular genetics. Eye (Lond) 31, 68–86 (2017). 10.1038/eye.2016.198

3 Heermann, S., Schutz, L., Lemke, S., Krieglstein, K. & Wittbrodt, J. Eye morphogenesis driven by epithelial flow into the optic cup facilitated by modulation of bone morphogenetic protein. Elife 4 (2015). 10.7554/eLife.05216

4 Kwan, K. M., Otsuna, H., Kidokoro, H., Carney, K. R., Saijoh, Y. & Chien, C. B. A complex choreography of cell movements shapes the vertebrate eye. Development 139, 359–372 (2012). 10.1242/dev.071407

5 Picker, A. et al. Dynamic coupling of pattern formation and morphogenesis in the developing vertebrate retina. PLoS Biol 7, e1000214 (2009). 10.1371/journal.pbio.1000214

6 Fitzpatrick, D. R. & van Heyningen, V. Developmental eye disorders. Curr Opin Genet Dev 15, 348–353 (2005). 10.1016/j.gde.2005.04.013

7 Gestri, G., Bazin-Lopez, N., Scholes, C. & Wilson, S. W. Cell Behaviors during Closure of the Choroid Fissure in the Developing Eye. Front Cell Neurosci 12, 42 (2018). 10.3389/fncel.2018.00042

8 James, A., Lee, C., Williams, A. M., Angileri, K., Lathrop, K. L. & Gross, J. M. The hyaloid vasculature facilitates basement membrane breakdown during choroid fissure closure in the zebrafish eye. Dev Biol 419, 262–272 (2016). 10.1016/j.ydbio.2016.09.008

9 Eckert, P., Knickmeyer, M. D. & Heermann, S. In Vivo Analysis of Optic Fissure Fusion in Zebrafish: Pioneer Cells, Basal Lamina, Hyaloid Vessels, and How Fissure Fusion is Affected by BMP. Int J Mol Sci 21 (2020). 10.3390/ijms21082760

10 Hardy, H. & Rainger, J. Cell adhesion marker expression dynamics during fusion of the optic fissure. Gene Expr Patterns 50, 119344 (2023). 10.1016/j.gep.2023.119344

11 Chang, L., Blain, D., Bertuzzi, S. & Brooks, B. P. Uveal coloboma: clinical and basic science update. Curr Opin Ophthalmol 17, 447–470 (2006). 10.1097/01.icu.0000243020.82380.f6

12 Gregory-Evans, C. Y., Williams, M. J., Halford, S. & Gregory-Evans, K. Ocular coloboma: a reassessment in the age of molecular neuroscience. J Med Genet 41, 881–891 (2004). 10.1136/jmg.2004.025494

13 AS, A. L., Gregory-Evans, C. Y. & Gregory-Evans, K. An update on the genetics of ocular coloboma. Hum Genet 138, 865–880 (2019). 10.1007/s00439-019-02019-3

14 Selzer, E. B., Blain, D., Hufnagel, R. B., Lupo, P. J., Mitchell, L. E. & Brooks, B. P. Review of evidence for environmental causes of uveal coloboma. Surv Ophthalmol 67, 1031–1047 (2022). 10.1016/j.survophthal.2021.12.008

15 Shah, S. P. et al. Anophthalmos, microphthalmos, and typical coloboma in the United Kingdom: a prospective study of incidence and risk. Invest Ophthalmol Vis Sci 52, 558–564 (2011). 10.1167/iovs.10-5263

16 Morrison, D. et al. National study of microphthalmia, anophthalmia, and coloboma (MAC) in Scotland: investigation of genetic aetiology. J Med Genet 39, 16–22 (2002). 10.1136/jmg.39.1.16

17 Hornby, S. J., Dandona, L., Jones, R. B., Stewart, H. & Gilbert, C. E. The familial contribution to non-syndromic ocular coloboma in south India. Br J Ophthalmol 87, 336–340 (2003). 10.1136/bjo.87.3.336

18 Patel, A. & Sowden, J. C. Genes and pathways in optic fissure closure. Semin Cell Dev Biol 91, 55–65 (2019). 10.1016/j.semcdb.2017.10.010

19 Harding, P. & Moosajee, M. The Molecular Basis of Human Anophthalmia and Microphthalmia. J Dev Biol 7 (2019). 10.3390/jdb7030016

20 Sudol, M. Yes-associated protein (YAP65) is a proline-rich phosphoprotein that binds to the SH3 domain of the Yes proto-oncogene product. Oncogene 9, 2145–2152 (1994).

21 Ma, S., Meng, Z., Chen, R. & Guan, K. L. The Hippo Pathway: Biology and Pathophysiology. Annu Rev Biochem 88, 577–604 (2019). 10.1146/annurev-biochem-013118-111829

22 Zhang, H. et al. TEAD transcription factors mediate the function of TAZ in cell growth and epithelial-mesenchymal transition. J Biol Chem 284, 13355–13362 (2009). 10.1074/jbc.M900843200

23 Zhao, B. et al. TEAD mediates YAP-dependent gene induction and growth control. Genes Dev 22, 1962–1971 (2008). 10.1101/gad.1664408

24 Williamson, K. A. et al. Heterozygous loss-of-function mutations in YAP1 cause both isolated and syndromic optic fissure closure defects. Am J Hum Genet 94, 295–302 (2014). 10.1016/j.ajhg.2014.01.001

25 DeYoung, C., Guan, B., Ullah, E., Blain, D., Hufnagel, R. B. & Brooks, B. P. De novo frameshift mutation in YAP1 associated with bilateral uveal coloboma and microphthalmia. Ophthalmic Genet 43, 513–517 (2022). 10.1080/13816810.2022.2028299

26 Oatts, J. T., Savar, L. & Hwang, D. G. Late extrusion of intrastromal corneal ring segments: A report of two cases. Am J Ophthalmol Case Rep 8, 67–70 (2017). 10.1016/j.ajoc.2017.10.004

27 Holt, R. et al. New variant and expression studies provide further insight into the genotype-phenotype correlation in YAP1-related developmental eye disorders. Sci Rep 7, 7975 (2017). 10.1038/s41598-017-08397-w

28 Oatts, J. T., Hull, S., Michaelides, M., Arno, G., Webster, A. R. & Moore, A. T. Novel heterozygous mutation in YAP1 in a family with isolated ocular colobomas. Ophthalmic Genet 38, 281–283 (2017). 10.1080/13816810.2016.1188122

29 Morin-Kensicki, E. M. et al. Defects in yolk sac vasculogenesis, chorioallantoic fusion, and embryonic axis elongation in mice with targeted disruption of Yap65. Mol Cell Biol 26, 77–87 (2006). 10.1128/MCB.26.1.77-87.2006

30 Kim, S. et al. Ocular phenotypic consequences of a single copy deletion of the Yap1 gene (Yap1 (+/-)) in mice. Mol Vis 25, 129–142 (2019).

31 Kim, J. Y. et al. Yap is essential for retinal progenitor cell cycle progression and RPE cell fate acquisition in the developing mouse eye. Dev Biol 419, 336–347 (2016). 10.1016/j.ydbio.2016.09.001

32 Kimelman, D., Smith, N. L., Lai, J. K. H. & Stainier, D. Y. Regulation of posterior body and epidermal morphogenesis in zebrafish by localized Yap1 and Wwtr1. Elife 6 (2017). 10.7554/eLife.31065

33 Miesfeld, J. B. et al. Yap and Taz regulate retinal pigment epithelial cell fate. Development 142, 3021–3032 (2015). 10.1242/dev.119008

34 Lai, J. K. H. et al. The Hippo pathway effector Wwtr1 regulates cardiac wall maturation in zebrafish. Development 145 (2018). 10.1242/dev.159210

35 Hardy, H. et al. Detailed analysis of chick optic fissure closure reveals Netrin-1 as an essential mediator of epithelial fusion. Elife 8 (2019). 10.7554/eLife.43877

36 Lupo, G. et al. Retinoic acid receptor signaling regulates choroid fissure closure through independent mechanisms in the ventral optic cup and periocular mesenchyme. Proc Natl Acad Sci U S A 108, 8698–8703 (2011). 10.1073/pnas.1103802108

37 Vinothkumar, S., Rastegar, S., Takamiya, M., Ertzer, R. & Strahle, U. Sequential and cooperative action of Fgfs and Shh in the zebrafish retina. Dev Biol 314, 200–214 (2008). 10.1016/j.ydbio.2007.11.034

38 Sousa-Ortega, A. et al. A Yap-dependent mechanoregulatory program sustains cell migration for embryo axis assembly. Nat Commun 14, 2804 (2023). 10.1038/s41467-023-38482-w

39 Williamson, K. A. & FitzPatrick, D. R. The genetic architecture of microphthalmia, anophthalmia and coloboma. Eur J Med Genet 57, 369–380 (2014). 10.1016/j.ejmg.2014.05.002

40 George, A., Lee, J., Liu, J., Kim, S. & Brooks, B. P. Zebrafish model of RERE syndrome recapitulates key ophthalmic defects that are rescued by small molecule inhibitor of shh signaling. Dev Dyn 252, 495–509 (2023). 10.1002/dvdy.561

41 Lee, J., Willer, J. R., Willer, G. B., Smith, K., Gregg, R. G. & Gross, J. M. Zebrafish blowout provides genetic evidence for Patched1-mediated negative regulation of Hedgehog signaling within the proximal optic vesicle of the vertebrate eye. Dev Biol 319, 10–22 (2008). 10.1016/j.ydbio.2008.03.035

42 Kobayashi, T., Yasuda, K. & Araki, M. Coordinated regulation of dorsal bone morphogenetic protein 4 and ventral Sonic hedgehog signaling specifies the dorso-ventral polarity in the optic vesicle and governs ocular morphogenesis through fibroblast growth factor 8 upregulation. Dev Growth Differ 52, 351–363 (2010). 10.1111/j.1440-169X.2010.01170.x

43 Bharti, K., Liu, W., Csermely, T., Bertuzzi, S. & Arnheiter, H. Alternative promoter use in eye development: the complex role and regulation of the transcription factor MITF. Development 135, 1169–1178 (2008). 10.1242/dev.014142

44 Bumsted, K. M. & Barnstable, C. J. Dorsal retinal pigment epithelium differentiates as neural retina in the microphthalmia (mi/mi) mouse. Invest Ophthalmol Vis Sci 41, 903–908 (2000).

45 Martinez-Morales, J. R., Signore, M., Acampora, D., Simeone, A. & Bovolenta, P. Otx genes are required for tissue specification in the developing eye. Development 128, 2019–2030 (2001). 10.1242/dev.128.11.2019

46 Martinez-Morales, J. R. et al. OTX2 activates the molecular network underlying retina pigment epithelium differentiation. J Biol Chem 278, 21721–21731 (2003). 10.1074/jbc.M301708200

47 Lister, J. A., Robertson, C. P., Lepage, T., Johnson, S. L. & Raible, D. W. nacre encodes a zebrafish microphthalmia-related protein that regulates neural-crest-derived pigment cell fate. Development 126, 3757–3767 (1999). 10.1242/dev.126.17.3757

48 Lister, J. A., Close, J. & Raible, D. W. Duplicate mitf genes in zebrafish: complementary expression and conservation of melanogenic potential. Dev Biol 237, 333–344 (2001). 10.1006/dbio.2001.0379

49 Sinagoga, K. L., Larimer-Picciani, A. M., George, S. M., Spencer, S. A., Lister, J. A. & Gross, J. M. Mitf-family transcription factor function is required within cranial neural crest cells to promote choroid fissure closure. Development 147 (2020). 10.1242/dev.187047

50 Lane, B. M. & Lister, J. A. Otx but not Mitf transcription factors are required for zebrafish retinal pigment epithelium development. PLoS One 7, e49357 (2012). 10.1371/journal.pone.0049357

51 Farhy, C. et al. Pax6 is required for normal cell-cycle exit and the differentiation kinetics of retinal progenitor cells. PLoS One 8, e76489 (2013). 10.1371/journal.pone.0076489

52 Bharti, K. et al. A regulatory loop involving PAX6, MITF, and WNT signaling controls retinal pigment epithelium development. PLoS Genet 8, e1002757 (2012). 10.1371/journal.pgen.1002757

53 Hitchcock, P. F., Macdonald, R. E., VanDeRyt, J. T. & Wilson, S. W. Antibodies against Pax6 immunostain amacrine and ganglion cells and neuronal progenitors, but not rod precursors, in the normal and regenerating retina of the goldfish. J Neurobiol 29, 399–413 (1996). 10.1002/(SICI)1097-4695(199603)29:3<399::AID-NEU10>3.0.CO;2-4

54 Halder, G., Dupont, S. & Piccolo, S. Transduction of mechanical and cytoskeletal cues by YAP and TAZ. Nat Rev Mol Cell Biol 13, 591–600 (2012). 10.1038/nrm3416

55 Gordon, H. B., Lusk, S., Carney, K. R., Wirick, E. O., Murray, B. F. & Kwan, K. M. Hedgehog signaling regulates cell motility and optic fissure and stalk formation during vertebrate eye morphogenesis. Development 145 (2018). 10.1242/dev.165068

56 Kimmel, C. B., Ballard, W. W., Kimmel, S. R., Ullmann, B. & Schilling, T. F. Stages of embryonic development of the zebrafish. Dev Dyn 203, 253–310 (1995). 10.1002/aja.1002030302

57 Sullivan-Brown, J., Bisher, M. E. & Burdine, R. D. Embedding, serial sectioning and staining of zebrafish embryos using JB-4 resin. Nat Protoc 6, 46–55 (2011). 10.1038/nprot.2010.165

